# Filament formation by the translation factor eIF2B regulates protein synthesis in starved cells

**DOI:** 10.1101/467829

**Authors:** Elisabeth Nüske, Guendalina Marini, Doris Richter, Weihua Leng, Aliona Bogdanova, Titus Franzmann, Gaia Pigino, Simon Alberti

## Abstract

Cells exposed to starvation have to adjust their metabolism to conserve energy and protect themselves. Protein synthesis is one of the major energy-consuming processes and as such has to be tightly controlled. The mechanism by which starved cells regulate the process of protein synthesis is largely unknown. Here, we report that the essential translation initiation factor eIF2B forms filaments in starved budding yeast cells. We demonstrate that filamentation is triggered by starvation-induced acidification of the cytosol, which is caused by an influx of protons from the extracellular environment. We show that filament assembly by eIF2B is necessary for rapid and efficient downregulation of translation. Importantly, this mechanism does not require the kinase Gcn2. Furthermore, analysis of site-specific variants of eIF2B suggests that eIF2B assembly results in enzymatically inactive filaments that promote stress survival and fast recovery of cells from starvation. We propose that translation regulation through protein assembly is a widespread mechanism that allows cells to adapt to fluctuating environments.

## Introduction

The ability of cells to effectively respond to stressful conditions is fundamental for their survival. Single-celled organisms such as *Saccharomyces cerevisiae* are frequently exposed to unfavorable environmental conditions such as starvation. Adaptation to stress conditions requires alterations in metabolism as well as the production of cytoprotective factors, such as molecular chaperones. One recently proposed survival strategy involves the formation of large protein assemblies. These assemblies are thought to protect proteins from damage (Franzmann et al., 2018), store proteins for later use (Franzmann et al., 2018; Laporte et al., 2008; Petrovska et al., 2014; Sagot et al., 2006) or down-regulate protein activity (Petrovska et al., 2014; Riback et al., 2017).

Glucose starvation induces re-localization of many cytoplasmic proteins into assemblies (Narayanaswamy et al., 2009; Noree et al., 2010). For unknown reasons, many of these assemblies adopt a highly regular filamentous structure. In the case of the two metabolic enzymes CTP synthase (CtpS) and glutamine synthetase (Gln1), filament formation has been shown to regulate enzymatic activity (Noree et al., 2014; Petrovska et al., 2014). However, the assembly mechanism and the function of most of these stress-induced filamentous assemblies remain unclear.

A major class of proteins that coalesce into cytoplasmic assemblies in starved cells are translation factors (Brengues and Parker, 2007; Franzmann et al., 2018; Hoyle et al., 2007; Noree et al., 2010). Protein synthesis is a cellular process that consumes a large amount of energy in growing cells. In fact, it has been estimated that this process can account for up to 50% of ATP consumption in eukaryotic cells (Hand and Hardewig, 1996). Thus, when energy is limited, for example upon glucose starvation or entry into stationary phase, cells must down-regulate translation to conserve energy and promote survival. Formation of cytoplasmic assemblies from translation factors in starved cells could be an adaptive strategy to regulate protein synthesis.

The process of protein synthesis is divided into three stages – initiation, elongation and termination – which all depend on a specific set of translation factors. Regulation of translation often occurs at the level of translation initiation (Hershey, 1991). For example, during amino acid starvation both the eukaryotic translation initiation factor 2 (eIF2) and its nucleotide exchange factor (GEF) eIF2B are targeted by signaling pathways that regulate their activity (Pavitt, 2005). eIF2 mediates the first step of translation initiation, where it binds the initiator methionyl-tRNA and forms a ternary complex that is involved in recognizing the start codon (Dever et al., 1995). Formation of this ternary complex only occurs when eIF2 is in its active GTP-bound state (Walton and Gill, 1975). eIF2-bound GTP is subsequently hydrolyzed to GDP at the ribosome and the active GTP-bound form of eIF2 is restored through a nucleotide exchange reaction that is mediated by eIF2B.

eIF2B is a decameric protein complex that consists of two heteropentamers. The protein subunits Gcd1 and Gcd6 form a catalytic subcomplex while Gcd2, Gcn3, and Gcd7 are components of a regulatory subcomplex. The eIF2B-catalyzed reaction is the rate-limiting step of translation initiation in stressed cells (reviewed in (Pavitt, 2005)). Under stress conditions, eIF2/eIF2B activity is regulated by post-translational modifications. In budding yeast, the kinase Gcn2 is the key player in this process. Gcn2 phosphorylates eIF2 and thus enhances the affinity of the initiation factor to its binding partner eIF2B. The tight binding of both initiation factors causes inhibition of the nucleotide exchange reaction and ultimately translational arrest (Krishnamoorthy et al., 2001). This reaction takes place in a variety of different stresses, such as amino acid starvation ((Hinnebusch and Fink, 1983), reviewed in (Simpson and Ashe, 2012)). Importantly, however, translational arrest during glucose starvation does not depend on Gcn2 (Ashe et al., 2000). Thus, alternative mechanisms must be in place to shut down translation during starvation, but these mechanisms have so far remained elusive.

Here, we show that the translation initiation factor eIF2B is diffusely distributed in exponentially growing yeast but re-localizes upon starvation, energy depletion and alcohol stress into multiple small assemblies that subsequently mature into filaments. We show that the trigger for filament formation is a stress-induced acidification of the cytosol and that filament assembly is necessary for rapid and efficient down-regulation of translation. Importantly, this mechanism is independent of the canonical stress signaling pathway mediated by the kinase Gcn2. We propose that eIF2B assembly into enzymatically inactive filaments is a protein-autonomous mechanism that allows yeast cells to rapidly adjust their protein synthesis rates to starvation conditions.

## Results

### eIF2B assembly formation is a response to starvation

The two essential translation initiation factors eIF2 and its GEF eIF2B can form mixed filamentous structures in budding yeast, but the mechanism of filament formation as well as the function has remained unclear (Campbell et al., 2005; Noree et al., 2010; Petrovska et al., 2014; Taylor et al., 2010). We hypothesized that filament formation by eIF2B could be an adaptation to energy depletion stress, because many proteins have been found to specifically form assemblies in starved cells or cells that enter into stationary phase (Franzmann et al., 2018; Munder et al., 2016; Petrovska et al., 2014; Rizzolo et al., 2017).

To investigate eIF2B filament formation in yeast cells, we employed live-cell fluorescence microscopy on cells expressing fluorescently labelled Gcn3 - the non-essential ***α***-subunit of eIF2B. To prevent tagging artifacts, a monomeric variant of superfolder GFP (sfGFP) was employed. Exponentially growing cells expressing Gcn3-sfGFP displayed no filaments; instead, the protein was evenly and diffusely distributed throughout the cytoplasm. However, in stationary phase, the natural form of nutrient scarcity, about 57% of the cells displayed filaments (Figure 1A). To investigate whether a lack of energy, which is characteristic for stationary phase cells, is causing filament formation, we subjected Gcn3-sfGFP expressing cells to two other forms of energy depletion: a milder form of energy depletion where we removed glucose from the medium and a more severe form of energy depletion where we additionally blocked glycolysis and mitochondrial respiration with chemicals (Dechant et al., 2010a). These treatments induced eIF2B filamentation in 8% and 86% of the cells, respectively (Figure 1B, Movie 1). This suggests that filament formation is a response to low energy levels and that the extent of filamentation depends on the magnitude of energy stress applied.

**Fig. 1.**
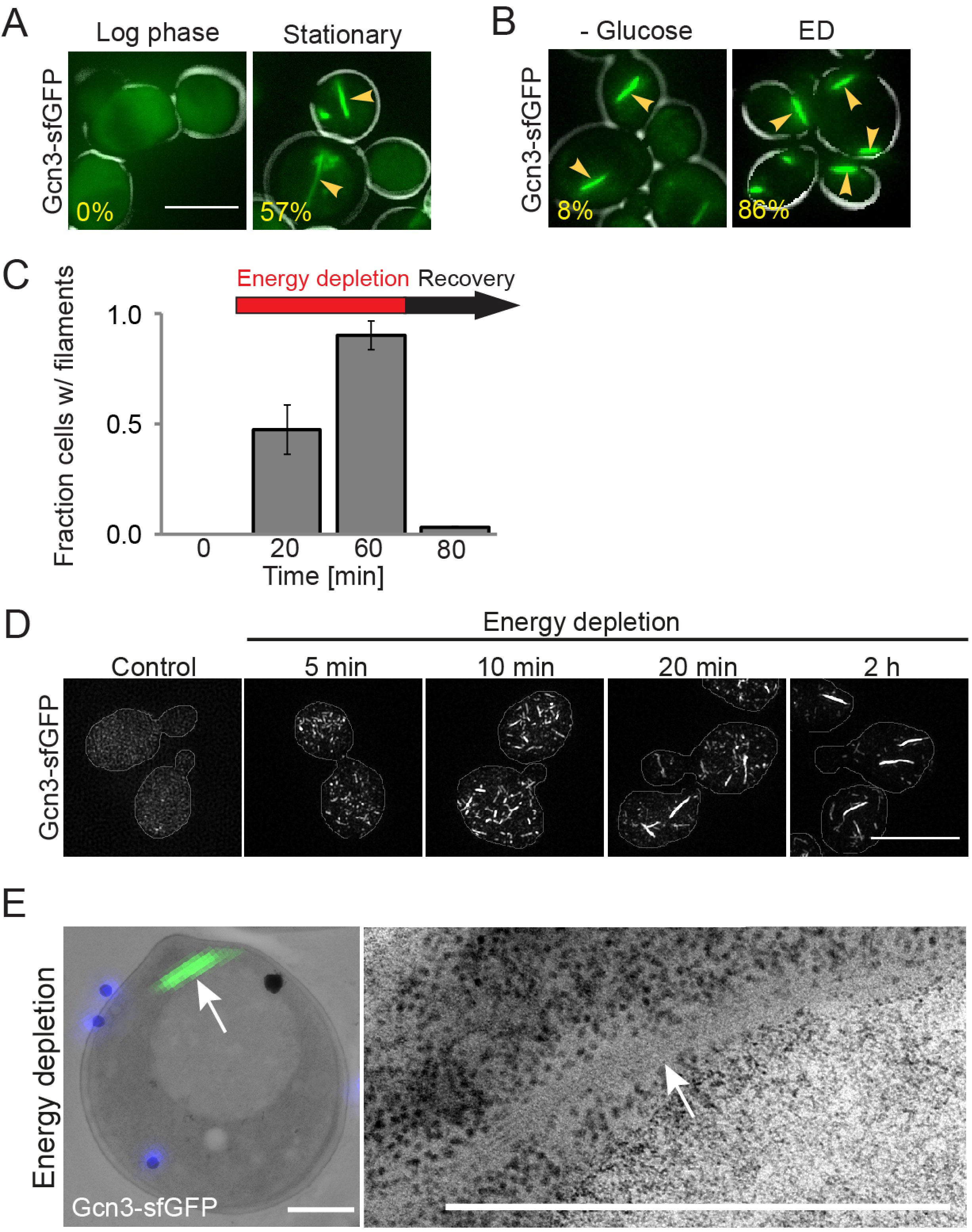
eIF2B filament formation is a starvation response. A) Live-cell fluorescence microscopy of *S. cerevisiae* expressing Gcn3-sfGFP(V206R) in log phase and after 3 days of growth in SC medium (Stationary). Note that log phase cells do not show filaments. Arrows point at filaments. The percentage of cells with filaments is shown at the lower left corner of each panel. Scale bar is 5 μm. B) Gcn3-sfGFP(V206R) localization after 30 min of glucose depletion (-Glucose) and after 30 min of energy depletion (ED) with 20 mM 2-deoxyglucose (2-DG), 10 jM antimycin (AM). Scale bar is 5 μm. C) Fraction of cells with Gcn3-sfGFP(V206R) filaments during different growth conditions, as quantified from live-cell fluorescence microscopy in a CellASICS microfluidic chamber. Cells grown to log phase in SC medium with 2% glucose (0 min), during ED (20 min and 60 min) and after recovery (80 min) are shown. The arrow indicates duration of ED (red) and recovery (black). D) Structured illumination microscopy of Gcn3-sfGFP(V206R) expressing log phase cells and after energy depletion. Scale bar is 5 μm. E) Ultrastructure of eIF2B filaments as found in energy depleted cells by correlative light and electron microscopy (CLEM). Left: electron micrograph of one representative cell overlaid with DAPI signal (blue – fiducials) and GFP signal (green – eIF2B filament). Right: close up of electron dense region corresponding to GFP signal. Scale bar is 1 μm.

Because eIF2B assemblies have also been described in unstressed and exponentially growing cells (Campbell et al., 2005), we tested whether the discrepancy with our data could be explained by the used fluorescent tag. Campbell et al. used a version of EGFP that is known to dimerize and reported that the abundance of eIF2B filaments depends on the position of the EGFP tag. To investigate whether the position of the fluorophore influences the result, we independently tagged all five eIF2B subunits with sfGFP as well as one of the eIF2 subunits (Sui2). However, in all cases filaments formed exclusively in energy-depleted cells and not in growing cells (Figure S1A). Moreover, tagging of Gcn3 with mCherry as well as with HA, a short epitope tag, produced identical results (Figure S1B and C). Taken together, these findings show that our fluorescent tag neither prevents nor promotes assembly formation and that tagged Gcn3 indicates the localization of the whole eIF2B complex. Most importantly these data demonstrate that eIF2B filaments form only in response to starvation but not in non-stressed cells.

### eIF2B assembly formation is not a result of protein misfolding and aggregation

Starvation stress challenges cellular proteostasis and often coincides with protein aggregation. This opens the possibility that eIF2B filaments are structures of misfolded and aggregated protein. To investigate this, we induced eIF2B filament formation by depleting cells of energy for one hour and then added back nutrients and imaged the cells during the recovery phase. We found that pre-formed assemblies dissolved within less than 10 minutes after release from stress (Figure 1C, Movie 2). Such short dissolution times are not indicative of misfolded and aggregated protein. Furthermore, the localization and disassembly kinetics were unaffected in the absence of the molecular protein disaggregase Hsp104, which is essential for clearing misfolded proteins during stress, demonstrating that the filaments are not a result of misfolding and aggregation (Figure S1D). In agreement with this, eIF2B assemblies were also sensitive to SDS and hence are not amyloidlike aggregates (Figure S1E). Altogether, these data show that eIF2B filaments are not protein aggregates, but protein assemblies that rapidly form and dissolve in response to starvation.

### eIF2B assembly involves bundling into filaments

We noticed in our fluorescence microscopy experiments that eIF2B not only formed filaments, but also multiple smaller assemblies in response to different starvation conditions. Live-cell time-lapse microscopy revealed that these assemblies form earlier than the filaments, suggesting that they are precursors of the larger filamentous structure (Movie 1). To further investigate this, we performed structured illumination microscopy (SIM) on cells that had been treated by energy depletion and were fixed at different time points after the onset of stress. This experiment revealed that multiple small round and comma-shaped assemblies had already formed after five minutes of energy depletion (Figure 1D). Shorter filaments were observed at later time points (10 minutes) and increased in length and intensity over time (20 minutes and 2 hours; Figure 1D), while the number of small assemblies decreased. This temporal pattern of assembly suggests that large filaments are in fact bundles of smaller filamentous structures. Similar bundles of filaments have previously been described for the metabolic enzyme Gln1 (Petrovska et al., 2014).

We further investigated the filament bundles of eIF2B by utilizing correlative light and electron microscopy (CLEM) on energy depleted cells expressing Gcn3-sfGFP. CLEM analysis revealed a regular pattern of several electron dense lines in the electron micrograph, which colocalized with the fluorescent signal of the eIF2B filaments and occupied a ribosome-free space (Figure 1E, also see accompanying manuscript by Marini et al.). We therefore conclude that eIF2B assemblies are bundles of individual filaments, which comprise most of the initially diffuse protein.

### eIF2B assembly is caused by starvation-induced cytosolic acidification

Our results so far show that the essential translation factor eIF2B forms filaments as a response to starvation and energy depletion. Moreover, filament formation is not the result of protein misfolding and aggregation but may be a physiological response to energy depletion stress. This raises questions about the cellular events regulating filament formation.

Starvation has previously been shown to be accompanied by a drop in cytosolic pH (pHc) and an increase in intracellular crowding (Joyner et al., 2016; Munder et al., 2016; Orij et al., 2009). Both parameters have strong effects on protein solubility and they are known driving forces for the formation of assemblies in starved cells. One striking example is provided by the metabolic enzyme Gln1, which forms filaments in a pH- and crowding-dependent manner (Petrovska et al., 2014). To test whether a reduced pHc is also required for eIF2B assembly formation, we determined if there is a temporal correlation between pH changes and eIF2B localization. Using time-dependent flow-cytometry analysis of energy-depleted cells expressing the pH-sensitive GFP variant pHluorin2 (Mahon, 2011a), we observed that the pHc, which we found to be around 7.6 in exponentially growing cells, dropped almost immediately after stress onset (Figure S2A). After only 3 minutes, the pHc adopted values around 6. This time frame is comparable to the onset of eIF2B assembly formation (Movie 1) and suggests, that eIF2B assembly and pH changes are linked.

To test whether the observed pHc reduction is necessary for assembly formation, we determined if eIF2B forms assemblies in energy-depleted cells exposed to neutral pH medium (a neutral pH of the medium prevents acidification of the cytoplasm, see (Munder et al., 2016)). This treatment completely prevented eIF2B assembly, while energy-depleted control cells in acidic medium still contained filaments (Figure 2A). To determine whether the reduced pHc is sufficient to induce filament formation, we altered the pHc directly by treating cells with a membrane-permeable protonophore in the presence of glucose. Filament formation was observed only in cells exposed to a low pH but not in cells exposed to a neutral pH (Figure 2A). These data show that cytosolic acidification is sufficient and required for filament formation by eIF2B.

**Fig. 2.**
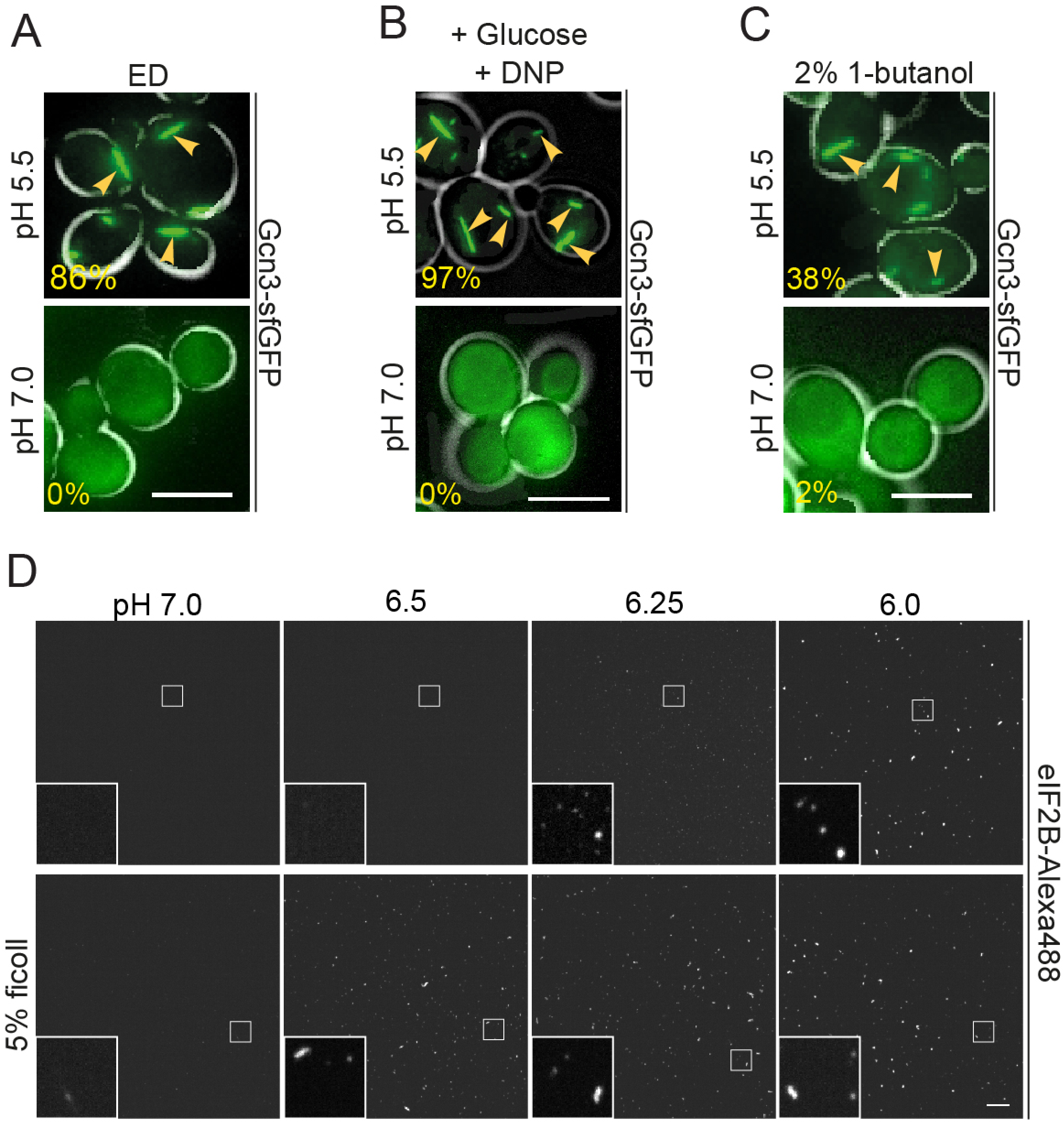
Cytosolic acidification triggers eIF2B filament formation. A) Live-cell imaging of Gcn3-sfGFP(V206R) filament formation after 20 min of ED at pH 5.5 (upper panel) or pH 7.0 (lower panel). The percentage of cells with filaments is shown in lower left corner of each panel. Note that no filaments form during ED at neutral pH. Scale bar is 5 μm. B) Imaging of cells after 20 min of treatment with DNP buffer with 2% glucose at pH 5.5 (upper panel) and pH 7.0 (lower panel). Scale bar is 5 μm. C) Imaging of cells after 20 min of treatment with 2% 1-butanol at pH 5.5 (upper panel) and pH 7.0 (lower panel). Note that no filaments form at neutral pH. Scale bar is 5 μm. D) Fluorescence microscopy of purified eIF2B after 10 min *in vitro* reconstitution. eIF2B-Alexa488 was mixed 1:20 with unlabeled eIF2B. Upper panels: The protein mix was incubated in pH 7.0, 6.5, 6.25, 6.0 (left to right) buffer at a final protein concentration of 0.1 μM. Lower panels: Same pH conditions as above but 5% ficoll was added. Scale bar is 10 μm.

Our findings so far show that eIF2B assembly is induced by cytosolic acidification. This suggests that eIF2B filaments could also form in response to other stress conditions that are known to alter the pHc. Interestingly, eIF2B assemblies have previously been described in cells treated with fusel alcohols such as 1-butanol (Taylor et al., 2010). Alcohol stress is known to alter the properties of the plasma membrane, thus making it more permeable to ions (Ingram, 1986). We therefore hypothesized that the effect of fusel alcohols on eIF2B may be mediated by an increased membrane permeability to protons and a subsequent acidification of the cytosol. To test this hypothesis, we determined whether our Gcn3-sfGFP strain forms filaments upon treatment with butanol. Indeed, cells treated with 2% butanol rapidly formed filaments (Figure 2C) and this treatment also led to a rapid drop in the pHc as measured with pHluorin2. The pHc quickly reversed to neutral values upon removal of butanol (Figure S2B) and at the same time filaments disassembled (data not shown). Importantly, filament formation was only observed in cells that were maintained in acidic medium, but not in cells maintained in neutral medium (Figure 2C). Based on these findings we conclude that filament formation by eIF2B is not limited to starvation, but rather takes place under conditions that induce a lower pHc. These data support the hypothesis that pHc changes may be a general signal for cellular stress, as previously proposed (Munder et al., 2016).

Is pH the only necessary factor to induce assembly formation by eIF2B? To test this, we purified eIF2B from insect cells and tested for the formation of assemblies in vitro. At a protein concentration of 0.1 μM, small assemblies formed at pH values of 6.25 and below (Figure 2D, S2D). These assemblies were reminiscent of assemblies seen at early time points during the exposure of cells to energy depletion conditions (Figure 1D). For unknown reasons, however, we did not observe larger filamentous structures. Thus, we speculate that bundling into larger filaments is dependent on additional factors that are missing from our in vitro reconstitution assay.

Because previous studies have demonstrated that crowding conditions affect the formation of various assemblies (Petrovska et al., 2014; Woodruff et al., 2017), we next tested the role of a polymeric crowder on eIF2B assembly. Indeed, assembly formation was promoted when we included 5% w/v ficoll in our *in vitro* assembly reaction (Figure 2D). In the presence of crowding agent, assemblies already formed at pH 6.5 as opposed to a pH 6.25 in the absence of crowder. In agreement with this, a related study found that glucose starvation and energy depletion also increase molecular crowding conditions inside cells (see accompanying manuscript by Marini et al.). To investigate whether changes in crowding conditions alone are sufficient to induce eIF2B assembly in cells, we exposed cells to different osmotic stress conditions. However, treatment with 1 M sorbitol or 0.6 M NaCl did not induce eIF2B assembly (Figure S2C). This indicates that pH changes are obligatory for assembly formation and that increased molecular crowding as it occurs during starvation, is not sufficient but may further enhance the effect of low pHc.

### eIF2B filament formation is necessary for efficient regulation of translation

Protein synthesis is the most energy-demanding processes in the cell and is down-regulated when cellular energy supply becomes limiting (Hand and Hardewig, 1996). We hypothesized that filament formation may be a cellular strategy to silence eIF2B activity and thus promote translational arrest during starvation. This makes the prediction that conditions that induce eIF2B assembly should also lead to translational arrest. Indeed, inhibition of translation initiation has previously been observed during glucose starvation and fusel alcohol stress (Ashe et al., 2001, 2000), conditions that also induce filament formation by eIF2B (Figure 1 and 2B). To investigate whether energy depletion also causes translational arrest, we performed polysome profiling analysis and a microscopy-based assay to measure translational activity in cells. For the latter, the methionine analogue homopropargylglycine (HPG) and a click-chemistry based approach were employed to determine the level of newly synthesized polypeptides. Indeed, energy depletion led to a rapid shutdown of translation and caused the disassembly of polysomes into monosomes (Figure 3A and B). Furthermore, ionophore-induced reduction of pHc caused a near-complete polysome runoff (Figure S3A). These data demonstrate that conditions that induce the formation of eIF2B filaments also cause translational arrest.

**Fig. 3.**
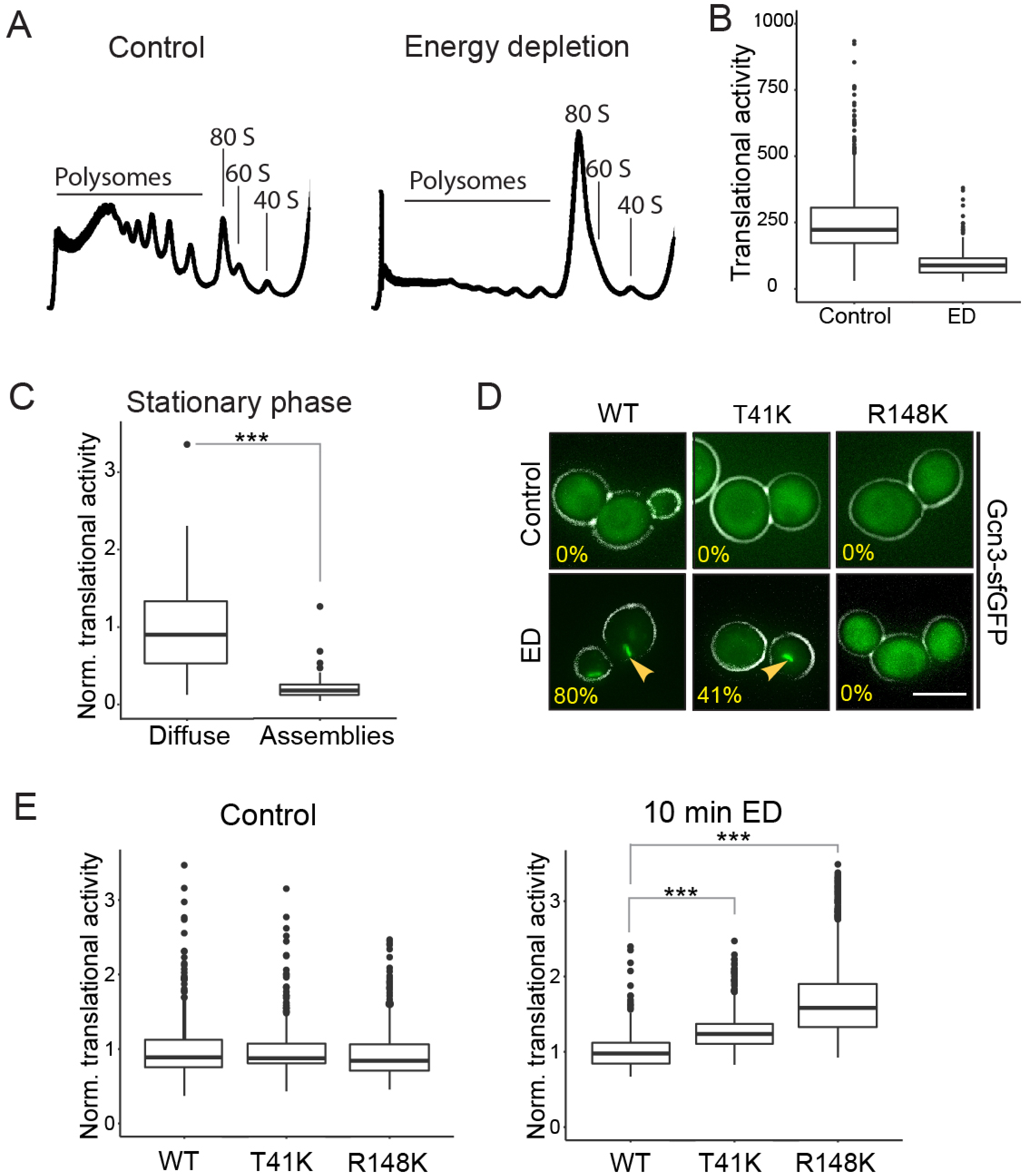
eIF2B filament formation facilitates down-regulation of translation. A) Polysome profiles of lysates from log phase cells (left) and after 10 min of energy depletion (right). Peaks corresponding to polysomes and to ribosomal subunits are indicated. Representative profiles from at least 3 independent experiments are shown. B) Translational activity as measured by HPG translation assay. The translational activity per cell is shown before and after 10 min of energy depletion (ED). Note, that translation is largely repressed within 10 min. C) Normalized translational activity of cells expressing Gcn3-sfGFP grown into stationary phase. Cells were grouped according to the localization of eIF2B. The translational activity of cells containing assemblies is shown on the right and the activity in cells with diffuse protein is shown on the left. Note that cells with assemblies exhibit significantly lower translational activity. ^⋆⋆⋆^ equals p<0.005 D) Fluorescence microscopy of wild-type Gcn3-sfGFP (WT) in comparison to Gcn3(T41K)-sfGFP (T41K) and Gcn3R148K-sfGFP (R148K) in control cells (upper panel) and after 10 min ED (lower panel). Percentage of cells with filaments is shown. Note that T41K shows fewer filaments and that filament formation is impaired in R148K. Scale bar is 5 μm. E) Normalized translational activity in WT cells and mutant cells under control conditions (left) and after 10 min of ED (right). Note that mutant cells continue translation longer during energy depletion. Values normalized to the average WT value from at least 3 independent experiments and >250 cells per condition are shown. ^⋆⋆⋆^ equals p<0.005

Interestingly, when determining the translation activity of stationary phase cells using the microscopy-based assay above, we noticed a large variation of translational activity from cell to cell. In agreement with this observation, we previously found that only ~57% of stationary phase cells had visible filaments (Figure 1B). Thus, we hypothesized that the translational activity of these cells depends on the extent of eIF2B filament formation. To test this, we grouped the stationary phase cells into cells with and without filaments and determined their translation activity. We found that cells with filaments showed significantly lower translation activity than cells without filaments (Figure 3C). These data indicate that filament formation and translation activity are correlated and suggest that filament formation may be required for the downregulation of translation.

Translational control on the level of eIF2B can be mediated by the kinase Gcn2. Stress conditions such as amino acid starvation trigger activation of Gcn2, which in turn phosphorylates eIF2, thus enhancing its binding to eIF2B and, ultimately, inhibiting nucleotide exchange (Krishnamoorthy et al., 2001). To test if eIF2B filament formation and down-regulation of translation under energy depletion conditions also depend on Gcn2, we monitored eIF2B localization before and after energy depletion in a Gcn2 deletion strain. eIF2B assembly was unaltered in the absence of Gcn2, indicating that filament assembly does not depend on eIF2 phosphorylation (Figure S3B). Furthermore, the polysome runoff in energy-depleted cells lacking Gcn2 was indistinguishable from the ones observed with wild-type cells (data not shown). These findings are in line with previous work showing that arrest of translation in response to glucose starvation is independent of Gcn2 (Ashe et al., 2000; Krishnamoorthy et al., 2001). These data suggest that the altered localization of eIF2B could be causative for the fast shut-down of translation during energy depletion.

To determine the function of filament formation, we created site-specific eIF2B variants, which exhibit impaired filament formation. These variants are based on a study by Taylor et al., who identified point mutations in Gcn3 (T41K and R148K) that decrease assembly formation of eIF2B (Taylor et al., 2010). We found that both variants formed less filaments during energy depletion (Figure 3D). The same trend was observed upon reduction of pHc to 6 (Figure S3C). While T41K was still able to assemble into filaments in some cells, no filamentation was observed in the R148K mutant (Figure 3D). Importantly, the mutations did not affect the protein expression level (Figure S3D).

We next tested whether a reduced ability to form filaments affects starvation-induced translational arrest. To this end, we determined the level of newly synthesized polypeptides within the first ten minutes of energy depletion (Figure 3E). These experiments revealed that both mutant strains continued translation longer than wild-type, indicating that filament formation promotes downregulation of translation. Furthermore, polysome profiling analysis showed faster polysome runoff in wild-type cells in comparison to both variants after ten minutes of energy depletion (Figure S3E). Taken together these data support the hypothesis that filament formation enables translation regulation upon stress by silencing eIF2B.

### eIF2B filament formation is essential for recovery and survival after starvation

Does the inability to form eIF2B filaments affect recovery from starvation stress? If eIF2B filament formation is a cellular strategy to down-regulate translation, cells that cannot form filaments may exhibit disadvantages during starvation. To test this, we first determined translation levels during recovery from energy starvation. We found that the two filamentation mutants showed defects in translation recovery. Thirty minutes after recovery from energy depletion, the translation level in R148K cells was nearly unchanged and T41K cells had recovered significantly less than wild-type cells (Figure 4A). These data suggest that efficient down-regulation of translation via filament formation is also important for re-starting translation after release from stress and re-growth.

**Fig. 4.**
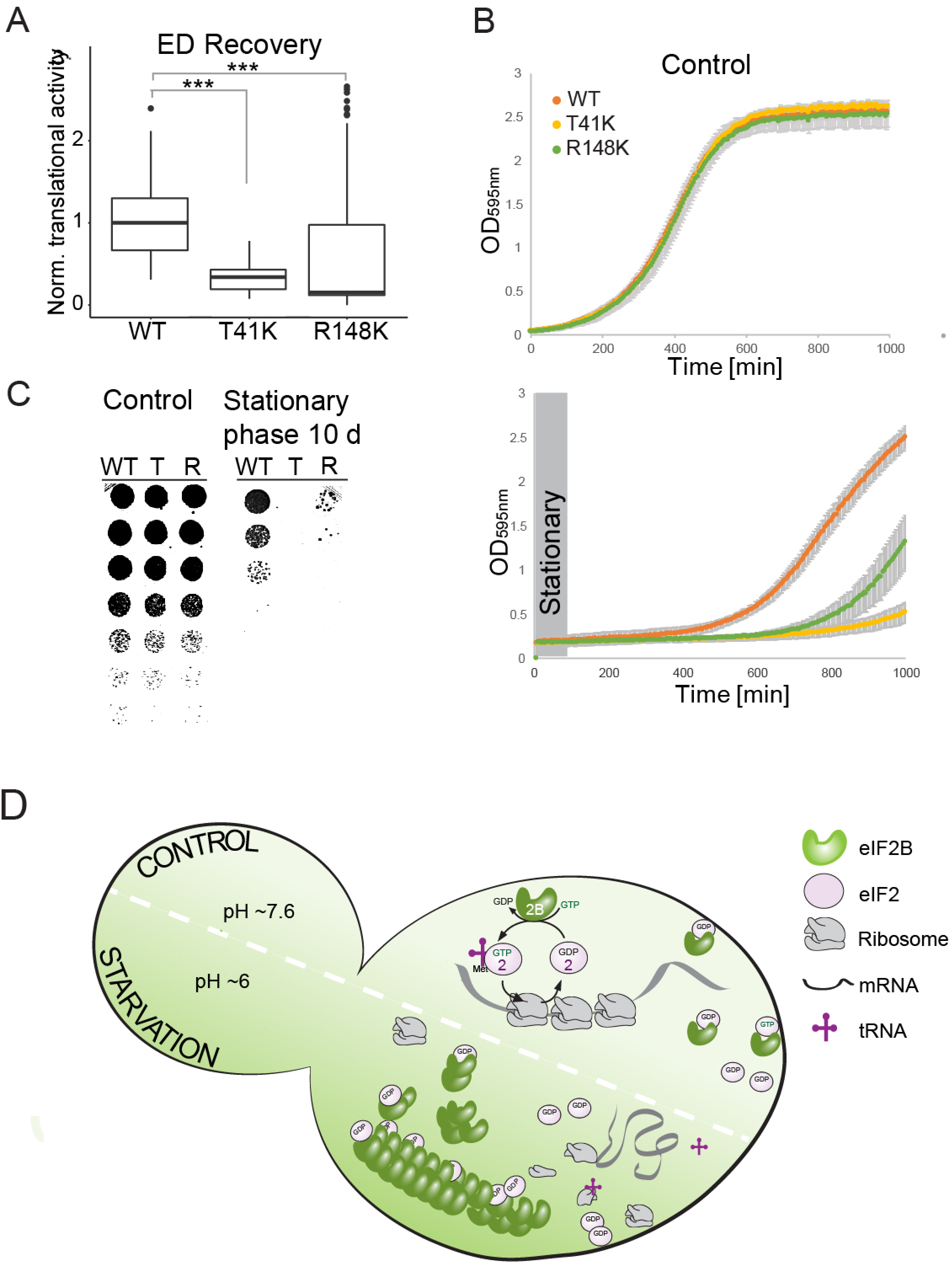
Filament formation is essential for recovery from and survival of starvation. A) Translational activity of cells expressing Gcn3, Gcn3(T41K) or Gcn3(R148K) after 20 min of recovery from 2 hours of energy depletion (ED Recovery) as measured by HPG translation assay. Note that wild-type cells show the highest level of translation after stress release. B) Growth curves of WT and mutant (T41K, R148K) strains diluted in SC medium to OD 0.05 after growth into log phase (upper panel) or stationary phase (lower panel). Growth was determined by measuring the optical density at 595 nm. Plots were generated from triplicates of 3 different biological replicates and the experiment was carried out at least 3 times. C) Spotting of five-fold serial dilutions of wild-type (WT), T41K (T) and R148K (R) cells grown into log phase (left) and after growth into stationary phase (right). D) Model for the molecular mechanism of starvation-induced filament formation of eIF2B and its impact on translation. The upper part of the cell represents the control situation, where the cytosolic pH is neutral and eIF2B is diffuse. Under these conditions the protein complex catalyzes the guanine nucleotide exchange for eIF2 which is essential for translation initiation. During starvation (lower part of the cell) the cytosolic pH is reduced to about 6, which triggers formation of eIF2B assemblies. Assembly formation silences eIF2B activity thus causing translation repression and possibly protection of the protein from aggregation.

Next, we tested how fast cell growth recovered from stationary phase or energy depletion. Both mutant strains exhibited a significantly longer lag phase compared to wild-type and thus recovery from energy starvation was slower (Figure 4B right, S4A). Importantly, all strains showed near-identical growth under non-stress control situation (Figure 4B left). To test if filament formation promotes cell survival during starvation, we grew cells into late stationary phase and tested their survival using a standard spotting assay. The survival assay revealed that T41K and R148K cells died faster than wild-type cells (Figure 4C). These findings are in agreement with the hypothesis that translation regulation via eIF2B filament formation is important for cellular adaptation to starvation and suggest that the process is important for cell survival.

## Discussion

In this paper, we show that the essential translation initiation factor eIF2B forms filaments upon a starvation-induced drop in cytosolic pH (Figure 4D). Acidification of the cytosol is triggered by low energy levels and accompanied by a reduction of the cytoplasmic volume (see accompanying manuscript by Marini et al.), which further increases the propensity of eIF2B to form filaments. Importantly, filament formation inactivates eIF2B and leads to translational arrest (Figure 4D). This translational arrest is required for cell survival under starvation conditions and promotes recovery from stress when cells are replenished with nutrients. Thus, eIF2B is the first confirmed translation initiation factor that forms adaptive assemblies under stress conditions, in addition to many other stress-induced assemblies that have recently been described to have adaptive roles (Franzmann et al., 2018; Munder et al., 2016; Petrovska et al., 2014; Riback et al., 2017; Rizzolo et al., 2017; Saad et al., 2017).

Numerous studies have shown that the kinase Gcn2 regulates translation through eIF2 phosphorylation, which enhances binding of eIF2 to eIF2B, thus inhibiting nucleotide exchange (Hinnebusch, 2005, 1993; Lindahl and Hinnebusch, 1992). Regulation of translation by Gcn2 has been demonstrated for diverse stresses such as heat stress or amino acid starvation (Farny et al., 2009; Mazroui et al., 2007; Pakos-Zebrucka et al., 2016). Surprisingly, however, we find that Gcn2 is not required for translational arrest under conditions in which cells are depleted of ATP. Similar findings have previously been made for glucose-depleted yeast cells (Ashe et al., 2000). This raises an important question: why do energy-depleted cells use a different mechanism for translational arrest?

We speculate that this has to do with the fact that eIF2B filamentation does not require energy in the form of ATP. Filament formation is induced by cytosolic acidification and occurs spontaneously in cells that are exposed to starvation conditions. By contrast, phosphorylation is an energetically costly process that uses substantial amounts of ATP and is limited under starvation conditions. Phosphate removal from eIF2 in the recovery phase allows the restoration of normal translation rates. In the case of translation regulation by filament formation, the filaments have to be disassembled when the stress subsides. This is achieved by removing protons from the cytosol to re-establish a neutral cytosolic pH. This means that energy is only consumed in the stress recovery phase when cells have enough nutrients to generate ATP. Thus, energy conservation may be one of the reasons why eIF2B and many proteins form higher-order assemblies in cells that are exposed to starvation conditions.

We find that filament formation by eIF2B is directly regulated by changes in cytosolic pH. Acidification of the cytosol is induced by removing nutrients such as glucose from the medium or by blocking glycolysis and mitochondrial respiration with drugs. All these conditions lower the cellular level of ATP and thus impair the removal of protons from the cytosol through ATP-dependent transporters and pumps (Dechant et al., 2010a; Munder et al., 2016; Orij et al., 2009). Importantly, we find that there are other conditions besides starvation that acidify the cytosol. For example, exposure of yeast cells to fusel alcohols such as butanol induces a rapid decrease in cytosolic pH (Figure S2B). We speculate that fusel alcohols change the permeability of the plasma membrane and thus cause a sudden influx of protons from the extracellular environment. There may be additional stresses that change the permeability of membranes and thus induce changes in cytosolic pH. For example, heat shock has been reported to cause a drop in cytosolic pH (Weitzel et al., 1987, 1985). This suggests that changes in cytosolic pH could be a more general signal to mount adaptive stress responses.

One previous study investigated the formation of filaments by eIF2B. The authors of this study proposed that filament formation increases the nucleotide exchange rate of eIF2B (Campbell et al., 2005). This was based on the observation that exponentially growing cells harbored eIF2B filaments. However, we could not detect eIF2B filaments in growing cells but found filaments exclusively in cells exposed to starvation conditions. This suggests that filament formation is a specific adaptation to conditions in which energy levels are low. Cells with low energy levels have to reduce their rate of translation, which can rapidly and most efficiently be achieved by decreasing eIF2B activity. In agreement with this, our data suggest that filament formation causes eIF2B inactivation. We attribute these discrepancies to the fact that different tags were used to visualize eIF2B. Indeed, Campbell *et al.* (2005) used a variant of GFP that is known to form weak dimers and this could have promoted the formation of filaments in exponentially growing cells through avidity effects (Landgraf et al., 2012). Thus, we conclude that previous occurrences of eIF2B filaments in growing cells could be the result of a tagging artifact.

Our data and those from other groups suggest that filament formation may occur via protomer stacking. We found that mutations in Gcn3 and Gcd1 drastically altered eIF2B filament formation, which indicates that these subunits are involved in stacking. This hypothesis is further supported by i) the crystal structure of eIF2B from *S. pombe* which shows that eIF2B dimerizes via its alpha subunit (Gcn3) (Kashiwagi et al., 2016), ii) work on human eIF2B and the regulatory subcomplex of *S. cerevisiae* eIF2B where a similar interaction of eIF2B-*α* was proposed (Bogorad et al., 2014; Wortham et al., 2014), iii) a study by Gordiyenko et al. who proposed dimerization of *S. cerevisiae* eIF2B pentamers via Gcd1 and Gcd6 – the catalytical subcomplex (Gordiyenko et al., 2014), and iv) data from Marini et al., who perform a comparison between the Cryo-EM structure of the eIF2B decamer and the in situ tomographic reconstruction of the eIF2B filament, which suggests a decamer-to-decamer interaction through the Gcd6 subunit. Altogether, this suggests that eIF2B stacking occurs via dimerization of the regulatory subcomplex on one side and the catalytic subcomplex on the other side. Our data and that of Marini et al. also reveal that eIF2B filaments are bundles of smaller filaments, which associate via lateral interactions. Such a mechanism has been described earlier for Gln1 filament formation (Petrovska et al., 2014) and may thus be of general relevance for other filament-forming proteins.

Our data with assembly-deficient eIF2B variants suggest that the nucleotide exchange activity of eIF2B is decreased by filament formation (Figure 3E). Currently, we do not know why filament formation leads to inactivation of eIF2B. We favor the view that eIF2B in its assembled state cannot undergo the conformational changes that are required to promote nucleotide exchange in eIF2. Indeed, guanine nucleotide exchange involves substantial structural rearrangements of eIF2 and eIF2B (Bogorad et al., 2017). Assembly of eIF2B into one large filament bundle likely prevents nucleotide exchange for another reason: the binding and release of eIF2 can no longer occur inside the bundle because of steric constraints. More detailed insight into the mechanism will require high resolution structures of assembled and unassembled eIF2B-eIF2 complexes. Unfortunately, we have not been able to perform direct measurements of the activity of assembled eIF2B, because we did not succeed in reconstituting filaments in vitro. Thus, future efforts should be directed towards finding conditions that allow the reconstitution of eIF2B filaments.

In growing cells, the intracellular pH is in the neutral range and thus eIF2B is fully active and translation occurs at a maximal rate. In response to starvation, lack of energy causes cytosolic acidification and assembly formation by eIF2B. This reduces the amount of active eIF2B and thus translation will increasingly come to a standstill. However, even starved cells need to keep a basal level of translation to maintain vital stress-protective processes. In agreement with this, we noticed that in starved cells not all the eIF2B signal is contained in filaments but that there is always a residual diffuse fluorescent signal. Interestingly, we also observed differences in the amount of assembled and unassembled eIF2B among cells. This may reflect the different abilities of cells to maintain a more neutral pH. The fine-tuning of filament formation via the cytosolic pH may give a population of yeast cells flexibility in responding to environmental changes. Thus, we speculate that eIF2B filament formation is a tunable system where the amount of assembled eIF2B is directly correlated with the cytosolic pH and thus the ability of a cell to keep its energy levels high.

We find that cells that cannot form filaments have specific defects in stress survival and recovery from stress. In agreement with this, cells that cannot form filaments show an impairment in shutting down translation. Starved yeast cells that continue to synthesize proteins at near-normal rates will presumably waste a lot of ATP and this could lead to a severe depletion of energy that eventually kills the cells. However, one can imagine that filaments have other functions than the ones described here. It is also possible that filament formation protects eIF2B from damage. This has previously been shown for the translation termination factor Sup35, which forms protective gels in cells exposed to energy depletion stress (Franzmann et al., 2018). Another possibility is that filament formation protects eIF2B from degradation and autophagy. Filaments may be spared from bulk autophagy, thus ensuring that a certain amount of eIF2B is available for restart of the cell cycle. The investigation of such alternative functions will require more work in the future.

In summary, we uncover a new mechanism of translation regulation in energy-starved cells. In the future, it will be interesting to investigate how pervasive this mechanism is. We suspect that organisms that live in fluctuating environments make widespread use of pH changes to regulate the material properties of their cytoplasm and form stress-adaptive assemblies that inactivate, protect and store proteins for later use.

## Materials and methods

### Yeast growth and strain generation

All yeast experiments were carried out in the genetic background of *S. cerevisiae* W303 *ADE*+ (*leu2*-*3112*; *his3*-*11*,-*15*; *trp1*-*1*; *ura3*-*1*; *can1*-*100*; [*PIN*+]). A list of all strains and plasmids used in this study can be found in Table 1. Cells were grown in yeast peptone dextrose (YPD), synthetic complete (SC) or synthetic dropout (SD) medium at 30°C unless stated differently. Yeast transformation was done using a standard lithium acetate/single-stranded carrier DNA/polyethylene glycol method (Daniel Gietz and Woods, 2002).

C-terminal tagging with sfGFP, mCherry and HA was carried out as described earlier (Sheff and Thorn, 2004). Positive clones were determined by microscopy and verified by Immunoblot analysis. Gene deletion was done according to the strategy described in (Gueldener et al., 2002). Positive clones were determined by sequencing of DNA amplified from single colonies. Integration of codon-optimized pHluorin2 (Mahon, 2011b) was done at the *trp* locus of W303 *ADE*+. Positive clones were identified via microscopy.

The filament formation mutant strains yMK16 and yMK54 described by Taylor et al. were tested positive for reduced filament formation during energy depletion. The genomic mutations in *GCN3* (T41K and R148K) and *GCD1* (P180S) from these strains were introduced into W303 *ADE*+ via the cloning free allele replacement strategy described by (Erdeniz et al., 1997). Positive clones were identified by sequencing of DNA amplified from single colonies. Because single point mutations in Gcn3 (T41K and R148K) alone caused a significant reduction of filament formation in the W303 *ADE*+ background we used, all following experiments were performed in the absence of the second mutation (Gcd1(P180S)).

### Cell stress

All stress treatments were carried out on cells grown to mid-log phase. For energy depletion, *S. cerevisiae* cells were washed twice and then incubated in SC medium without glucose but containing 20 mM 2-deoxyglucose (2-DG; Carl Roth GmbH, Karlsruhe, Germany) for inhibition of glycolysis, and 10 mM antimycin A for inhibition of mitochondrial respiration (Sigma-Aldrich, Steinheim, Germany). This treatment reduces intracellular ATP levels by more than 95% (Serrano, 1977). The intracellular pH of *S. cerevisiae* cells was adjusted by incubation in phosphate buffers of different pH in the presence of 2 mM 2,4-dinitrophenol (DNP; Sigma-Aldrich, Saint Louis, USA) as described previously (Dechant et al., 2010b; Munder et al., 2016; Petrovska et al., 2014). DNP buffers further contained 2% glucose. Stationary phase experiments were done by culturing cells for 2 days (starting from mid-log phase) in SC medium with 2% glucose at 30°C.

### Light microscopy

Samples for wide-field microscopy where either imaged in concanavalin-A-coated 4-well Matek dishes or in a CellASIC (Millipore) microfluidics flow setup combined with CellASIC ONIX Y04C microfluidic plates (Millipore). General fluorescence microscopy images of living or fixed cells and time-lapse movies were acquired using a Deltavision microscope system with softWoRx 4.1.2 software (Applied Precision). The system was based on an Olympus IX71 microscope, which was used with a 100× 1.4 NA UPlanSApo oil immersion objective. The images were collected with a CoolSnap HQ2 camera (Photometrics) at 1024×1024 (or 512×512) pixel files using 1×1 (or 2×2) binning. Images were maximum intensity projections of at least 12 individual images. Cell boundaries if shown (indicated by the white outline in fluorescence microscopy images) were introduced by changing the contrast of the DIC image and overlaying it with the fluorescent image.

### Structured illumination microscopy (SIM)

Stressed and control cells were fixed for at least 30 minutes in 3.7% formaldehyde. Glass slides were coated with concanavalin A for cell attachment and cells were immersed in Vectashield mounting medium prior imaging. SIM was carried out with a DeltaVision OMX v4 BLAZE (Applied Precision) microscope on an inverted stand, a 4x sCMOS camera (Andor) and a 100× 1.4 oil immersion UPlanSApochromat objective (Olympus). API softworx was used for driving the microscope and OMX SI reconstruction was carried out using Centos 4, the OMX SI reconstruction linux box. A 488 nm DPSS laser and a 561 nm DPSS laser were used to image GFP and mCherry signal respectively. At least five different Z-stacks (maximum Z-resolution) were recorded for each condition. Cell boundaries if shown (indicated by the white outline in fluorescence microscopy images) were drawn manually.

### Correlated light and electron microscopy (CLEM)

CLEM of yeast cells was done essentially as described in (Petrovska et al., 2014). For transmission electron microscopy (TEM), yeast cells were grown to log phase, vacuum filtered, mixed with 20% BSA, high pressure frozen (EMPACT2, Leica Microsystems, Wetzlar, Germany) and freeze-substituted with 0.1% uranyl acetate and 4% water in acetone at –90°C. Samples were transitioned into ethanol at –45°C, before infiltration into a Lowicryl HM-20 resin (Polysciences, Inc., Eppelheim, Germany), followed by UV polymerization at –25°C. Semi-thin (70,100,150 nm) sections were mounted on formvar-coated EM finder grids and stained for 3 min with lead citrate. Imaging was done in a Tecnai-12 biotwin TEM (FEI Company, Thermo Fisher Scientific) at 100 kV with a TVIPS 2k CCD camera (TVIPS GmbH, Gauting, Germany).

For CLEM, yeast cells expressing wild type or R23E Gcn3-sfGFP were processed for TEM. To allow alignment of EM and light microscopy (LM) images, unstained sections on EM grids were incubated with quenched 200 nm Blue (365/415) FluoSpheres as fiducials. Grids were mounted on a glass slide with VectaShield (Vector Laboratories, Inc., Burlingame, USA) and viewed in both green (for GFP) and UV (for the fiducials) channels. After staining for TEM, regions of interest in LM were relocated in TEM. Montaged images were acquired at multiple magnifications to facilitate the correlation. LM and TEM images were overlaid in ZIBAmira (Zuse-Institut, Berlin, Germany).

### pH measurement

For measurements of the cytosolic pH a codon-optimized version of the ratiometric fluorescent protein pHluorin2 was used. pH measurements and calculation during fusel alcohol stress were essentially done using an imaging-based method as described in (Munder et al., 2016). For pH measurement of individual cells, we use flow cytometry. The data sets were acquired using a FACS Aria Illu (Becton Dickinson) instrument with BV510/FITC and FITC/FITC filter sets. For all measurements, a single-cell gate was defined using the FSC-W and SSC-W parameters and only cells positive for pHluorin2 were considered. Gating was done using FlowJo software (FlowJo LLC, Ashland, OR, USA) and the relevant datasets where exported to Microsoft Excel or RStudio for further calculations. For pH calculation, a calibration curve for a pH range between pH 5 and pH 8 was determined using the protocol of (Diakov et al., 2013). After background subtraction, the mean emission ratio was calculated from the intensity readouts of both channels and plotted against pH to obtain a calibration curve. Subsequent pH measurements were calculated from a sigmoidal fit to the calibration curve. pH curves were plotted using R studio software. The results from kinetic pH measurements during energy depletion were fitted using a nonlinear asymmetric sigmoidal curve.

### Protein purification

For purification of His-tagged eIF2B, 500 ml of SF9 ESF cells (1×10^6^ cells/ml) were co-transfected 1:500 with recombinant baculovirus stocks for co-expression of eIF2B regulatory and catalytic subcomplexes (for plasmid names look at table 2). Cells were cultured for about 48 h and then harvested by mild centrifugation at 500 ×g for 5 min. Cells were resuspended in ice-cold lysis buffer (0.1 M Tris pH 8, 0.1 M KCl, 5 mM MgCl, 10% glycerol, 0.1% Triton-100, 0.02 M imidazole, protease inhibitors, 1 mM DTT) and lysed using an EmulsiFlex-C5 (Avestin). Cell lysates were clarified by centrifugation at 75,000 ×g for 30 minutes (rotor: JA 25.50, Beckman Coulter). The supernatant was filtered and loaded onto a 5 ml HisTrap column (GE Healthcare, Uppsala, Sweden). After washing with 75 ml of wash buffer (0.1 M Tris pH 8, 0.1 M KCl, 5 mM MgCl, 0.02 M imidazole, 1 mM DTT), the protein was eluted in elution buffer (wash buffer with 0.25 M imidazole) as 2.5 ml fractions. The fractions containing the protein were determined via Bradford assay and peak fractions were pooled prior to size-exclusion chromatography (SEC). SEC was performed using a Äkta Pure chromatography setup (GE Healthcare, Uppsala, Sweden), a Superose 6 Increase 5/150 GL column (GE Healthcare, Uppsala, Sweden) and buffer containing 0.05 M Tris pH 8, 0.15 M KCl and 2 mM MgCl. Purity of the protein was determined using standard SDS-PAGE and Coomassie Brilliant Blue staining protocols.

### In vitro reconstitution

Purified eIF2B was immediately used for reconstitution experiments. Some of the protein was labeled with Alexa 488 and the mixed 1:20 with unlabeled protein. The protein mix was then diluted in reconstitution buffers (0.1 M phosphate buffer pH 5.5-7.5, 0.15 M KCl, 2 mM MgCl, 1 mM DTT) to a final protein concentration of 0.33 μM. Two separate solutions containing 0.33 μM eIF2B in reconstitution buffers and 1.25%–10% PEG-20000 or 1.25%-15% ficoll 70 in reconstitution buffer were dispensed into two rows of a 96 well plate. A pipetting robot (Tecan, Model Freedom EVO 200) equipped with a TeMo 96 pipetting head with disposable tips (50 μl) was used to mix the protein (7.5 μl) and crowding solutions (17.5 μl) and then to transfer each condition in quadruplicate to a 384 well plastic bottom imaging plate (5 μl per well). The 384 well plate was imaged using a Yokogawa CV7000 high-content spinning disk confocal microscope and a 60x 1.2 NA water immersion objective. Z stacks (3 planes, 0.2 μm spacing) were taken at 4 different positions per well, once in solution and once close to the bottom surface. Z-stacks are presented as maximum intensity projections. Images shown represent pH 5.5-7.5 buffers +/- 5% ficoll.

### Yeast cell lysis

Cells were grown overnight in YPD medium to mid-log phase and then harvested by centrifugation at 3,000 rpm for 4 min. Cell pellets were washed once in dH_2_O and then resuspended in 500 μl cold lysis buffer (50 mM Tris pH 7.5; 150 mM NaCl; 2.5 mM EDTA; 1% (v/v) Triton X-100; 0.4 mM PMSF; 8 mM NEM; 1.25 mM benzamidine; 10 μg/ml pepstatin; 10 μg/ml chymostatin; 10 μg/ml aprotinin; 10 μg/ml leupeptin; 10 μg/ml E-64; protease inhibitors from Sigma-Aldrich, Steinheim, Germany) and added to about 500 μl ice-cold glass beads. Cells were lysed using mechanical disruption (bead beating) (Tissue Lyser II, Qiagen) at 25 Hz for 20-30 minutes. Unwanted cell debris and beads were removed by centrifugation at 8,000x for 10 minutes. Yeast cell lysates were subsequently used for immunoblotting, polysome profiling analysis or SDD-AGE.

### Immunostaining

For immunostaining, cells expressing Gcn3-HA were fixed via treatment with 3.7% formaldehyde (EMS, Hatfield, USA) for at least 30 min followed by 45 min incubation in spheroplasting buffer (100 mM phosphate buffer pH 7.5, 5 mM EDTA, 1.2 M Sorbitol (Sigma-Aldrich, Steinheim, Germany), Zymolyase (Zymo Research, USA)) at 30°C with mild agitation. Spheroplasts were permeabilized with 1% triton X-100 (Serva, Heidelberg, Germany), washed and then incubated with *α*-HA primary antibody from mouse (1:2000; Covance, USA) and Alexa Fluor 488 F(ab’)_2_ fragment of rabbit anti-mouse IgG (H+L) (1:2000; Invitrogen, A21204). GFP-tagged proteins were detected in lysates via western blot analysis using an *α*-GFP primary antibody (1:2000; Roche, Mannheim, Germany) and a secondary *α*-mouse antibody (1:5000; Sigma, Saint Louis, USA). The protein levels of PGK were determined as internal loading control with *α*-PGK primary antibody from mouse (1:5000; Invitrogen, Camarillo, USA) and secondary *α*-mouse antibody (1:5000; Sigma, Saint Louis, USA).

### SDD-AGE

Semi-denaturing detergent agarose gel electrophoresis (SDD-AGE) with lysates from filament-containing cells and control cells was performed essentially as described previously (Alberti et al., 2010). The supernatants of cell lysates were adjusted for equal protein concentrations and mixed 4:1 with 4x Sample buffer (40 mM Tris acetic acid, 2 mM EDTA, 20% glycerol, 4% SDS, bromophenol blue). Samples were incubated for 10 min at room temperature and loaded onto a 1.5% agarose gel containing 0.1% SDS in 1× TAE/0.1% SDS running buffer. The gel was run at 80 V. Proteins in the gel were detected by immunoblotting with a mCherry-specific antibody (MPI-CBG, Antibody facility).

### Translation assays

For polysome profiling, we used a protocol adapted from (Mašek et al., 2010). In brief, stressed and non-stressed cells were incubated with 0.1 mg/ml cycloheximide (AppliChem, Darmstadt, Germany), chilled and washed before cell lysis. Lysis was performed as described above but in buffer containing 0.1 mg/ml cycloheximide. Lysates were cleared by 10 min centrifugation at 8000 xg. Supernatant equivalent to about 20-25 OD_260nm_ were carefully layered on top of ice-cold sucrose gradients (7.5-50%) in ultra-clear centrifugation tubes (14×95 mm; Beckman Coulter, Brea, USA). After 2.5 h of ultra-centrifugation at 215,000 ×g in a SW40 Ti rotor (Beckman Coulter) at 4°C, a thin glass needle was placed at the bottom of the tube and gradients were gradually pump out (1 ml/min) using a peristaltic pump, while detecting the 260 nm signal using a BioCad LC 60 chromatography system (Applied Biosystems).

Translational activity was determined utilizing the Click-iT HPG Alexa 594 Fluor Protein Synthesis Assay Kit from Invitrogen (Molecular Probes, Eugene, USA). For control samples and energy depletion measurements, cells were incubated in SD - Met containing 50 μM of the methionine-analogue HPG. For stationary phase samples, HPG was added directly. After 30 min HPG incubation, the cells were fixed for at least 30 min (1 h for stationary phase) and then incubated for 45 min (90 min for stationary phase) in spheroplasting buffer at 30°C as described above. Spheroplasts were attached to clean, poly-L-lysine (Sigma-Aldrich, Steinheim, Germany)-coated Cell-View cell culture 4-well dishes (Greiner, Frickenhausen, Germany), washed, permeabilized with 1% triton X-100 and subsequently stained via click chemistry as described in the manufacturer’s protocol. All samples were thoroughly washed prior to imaging to minimize background signal. Imaging was carried out as described above using Cherry to detect the HPG signal. The translational activity per cell was determined from maximum intensity projections using Fiji by subtracting the background signal, manually selecting the area corresponding to each spheroplast in the field of view and reading out the mean intensity values for each area. All values of one experiment were then normalized to the average intensity of the wild-type sample. The combined data of all repeats were plotted as combination of box and dot plots using R Studio software. Statistics were calculated using a student’s paired t-test, with a two-tailed distribution.

### Growth and survival

*S. cerevisiae* wild type strain W303, were grown overnight, diluted to OD600 ~0.1 the next morning and regrown to OD600 ~0.5. Cells were harvested and resuspended in either phosphate buffers of pH 5.6-7.6, containing 2 mM DNP or in S-medium without glucose containing 20 mM 2-DG and 10 mM antimycin A. Cells were then incubated under shaking at 25°C. Samples were taken after 2, 24 and 48 h or as indicated in the specific experiments. Cells were washed once with H_2_O and subsequently spotted on YPD as five-fold serial dilutions.

For stress recovery growth assays, cells were grown to mid-log phase and energy depleted for 6 h or grown into stationary phase for 3 days. Control cells were diluted and regrown to OD ~0.5. Control cells and starved cells were diluted to OD ~0.05 in SC and added as triplicates to a 96 well plate. The plate was then incubated at 30°C for 200 cycles of 5 min in a FLUOstar Omega plate reader (BMG Labtech), OD_595_ was recorded with 10 flashes per well per cycle.

## Figure legends

**Supplementary Fig. 1.**
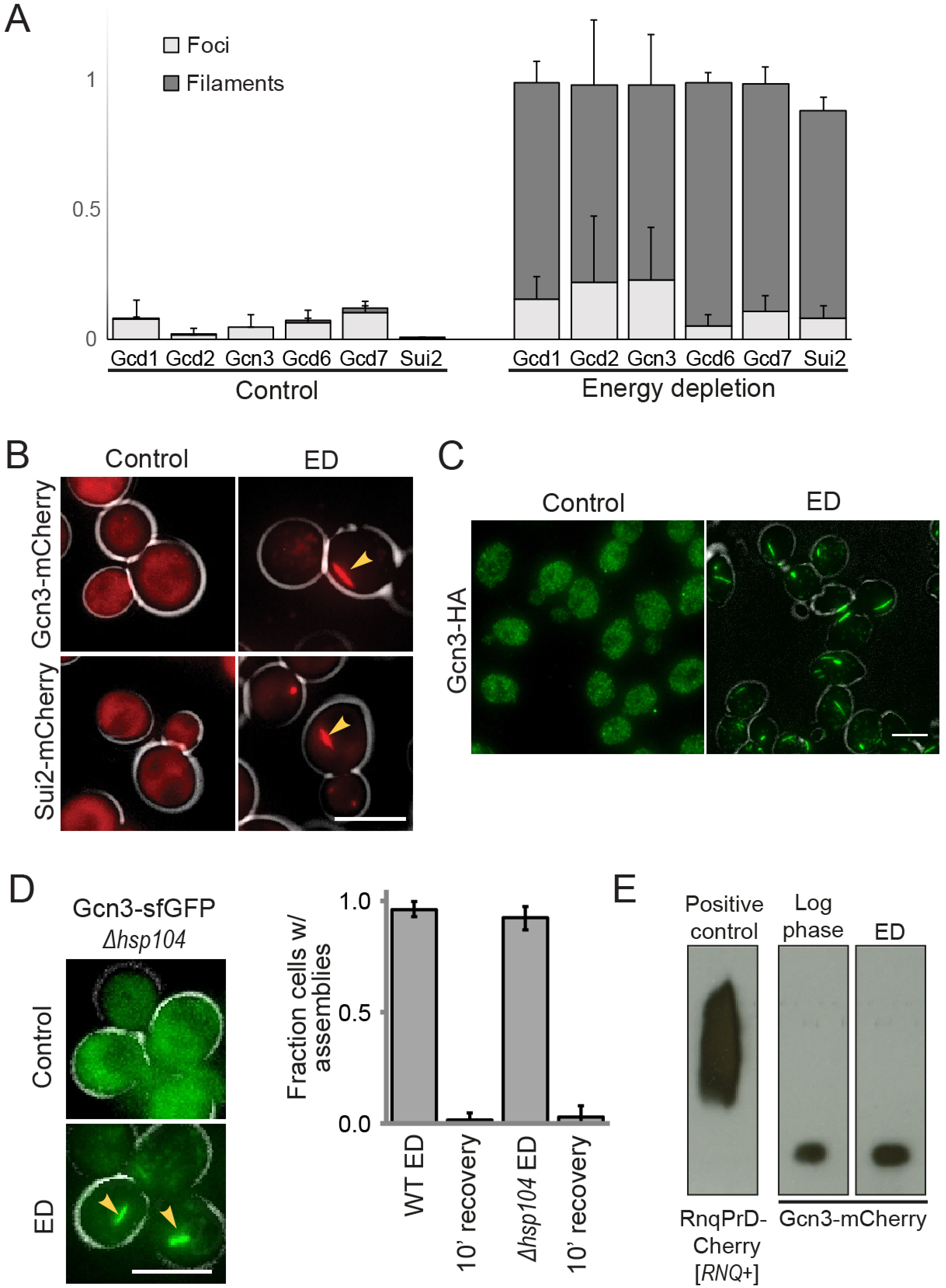
Filaments are starvation-induced protein assemblies. A) Fraction of cells with eIF2B assemblies (foci – light grey; filaments – dark grey) quantified from live-cell fluorescence microscopy images. Quantification was done for cells expressing subunits of eIF2B (Gcd1, Gcd2, Gcn3, Gcd6, Gcd7) and eIF2- (Sui2) tagged with sfGFP(V206R) during logarithmic growth (control, left) and after 20 min of energy depletion (ED, right). Results were obtained from at least 200 cells per sample and 3 independent experiments. B) Sui2-mCherry and Gcn3-mCherry localization in control cells and after 20 min of ED. Arrows point at filaments. C) Immunofluorescence of Gcn3-HA in fixed control cells and after 20 min of ED. Note that the protein is diffuse in control cells and localizes in filaments in starved cells. D) Live-cell fluorescence microscopy in *HSP104* deletion background (left) and quantification of eIF2B assemblies in Δ*hsp104* and wild type cells after 1 hour of ED and 10 min after recovery in SC medium. Note that deletion of *HSP104* does not affect filament formation or dissolution. Scale bar is 5 μm. E) SDD-AGE of cell lysates from Gcn3-mCherry expressing cells from log phase (middle) and after 1 hour of ED (right). Lysates from RnqPrD-Cherry [*RNQ*+] expressing cells were used as positive control (left).

**Supplementary Fig. 2.**
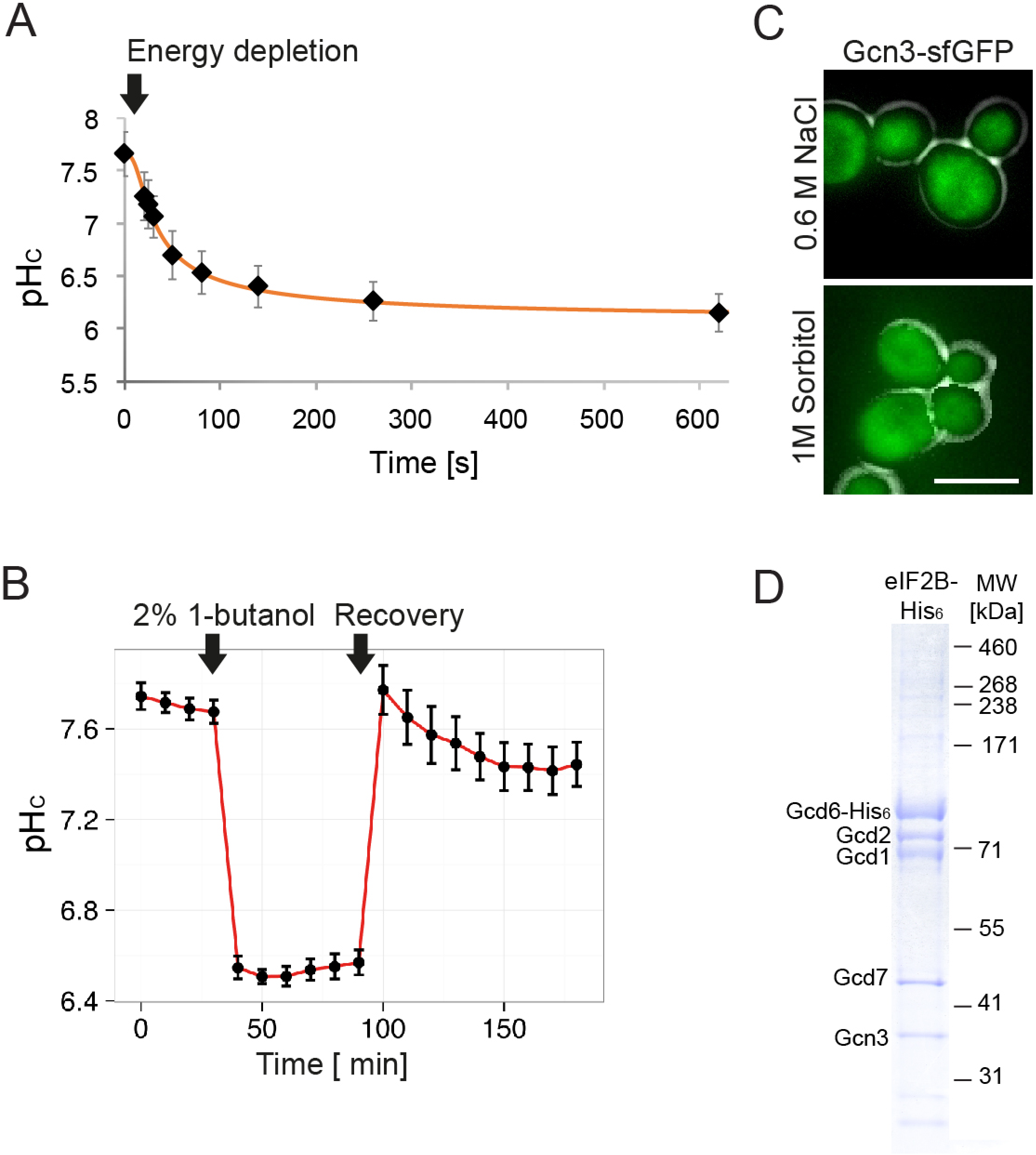
Filamentation-inducing conditions cause reversible acidification of the cytosol. A) pH_c_ changes in pHluorin2-expressing culture during energy depletion (ED) as measured by flow cytometry. Note that pH_c_ drops by one unit within 3 min. B) pH measurement during fusel alcohol stress as determined by live-cell fluorescence microscopy of yeast expressing pHluorin2. Note that pH_c_ drops rapidly upon addition of 2% 1-butanol and recovers to normal levels upon stress release. C) Fluorescence microscopy of Gcn3-sfGFP(V206R) expressing yeast after 20 min of treatment with 0.6 M NaCl (upper panel) and 1 M sorbitol (lower panel). Note that osmotic stress does not induce assembly formation. Scale bar is 5 μm. D) Acrylamide gel of eIF2B with His-tag after purification from insect cells. Protein bands are visualized via staining with Coomassie Brilliant Blue.

**Supplementary Fig. 3.**
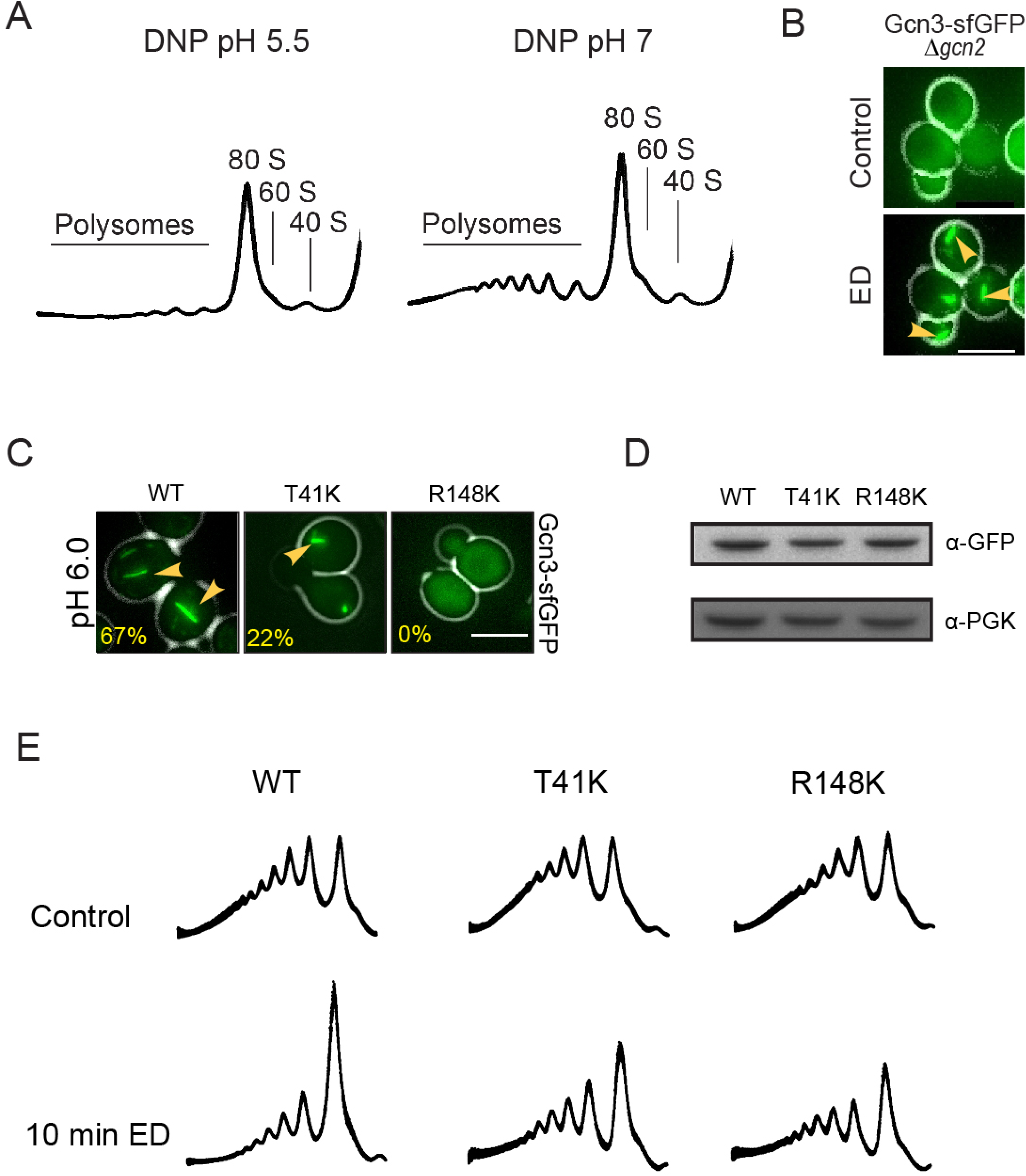
Filament formation is required for efficient translation downregulation. A) Polysome profiles after 10 min of treatment with 20 mM DNP pH 5.5 (left) and pH 7.0 (right) in the presence of 2% glucose (w/v). The regions corresponding to polysomes as well as the monosome subunits are indicated. B) Live-cell fluorescence microscopy of Gcn3-sfGFP-expressing cells lacking Gcn2. Control cells are shown in the upper panel and cells treated for 20 min with energy depletion (ED) are shown below. Note that Gcn2 deletion does not affect filament formation. Scale bar is 5 μm. C) Imaging of wild-type cells (WT) and mutant cells that showed filament formation deficiency during ED (T41K, R148K). Imaging was done after 20 min incubation in pH 6 buffer containing 2% glucose and 20 mM DNP. Note that T51K forms fewer filaments and no filaments form in R148K expressing cells. Scale bar is 5 μm. D) Western Blot of GFP-tagged Gcn3 from lysates of log phase cells expressing wild-type Gcn3, T41K or R148K. PGK1 was used as loading control. A GFP-specific antibody and a PGK-specific antibody were used for immunodetection, respectively. Note that WT, T41K and R148K protein level do not differ. E) Polysome profiles of WT, T41K and R148K lysates from cells grown to log phase (control), treated for 10 min by ED and after growth into stationary phase.

**Supplementary Fig. 4.**
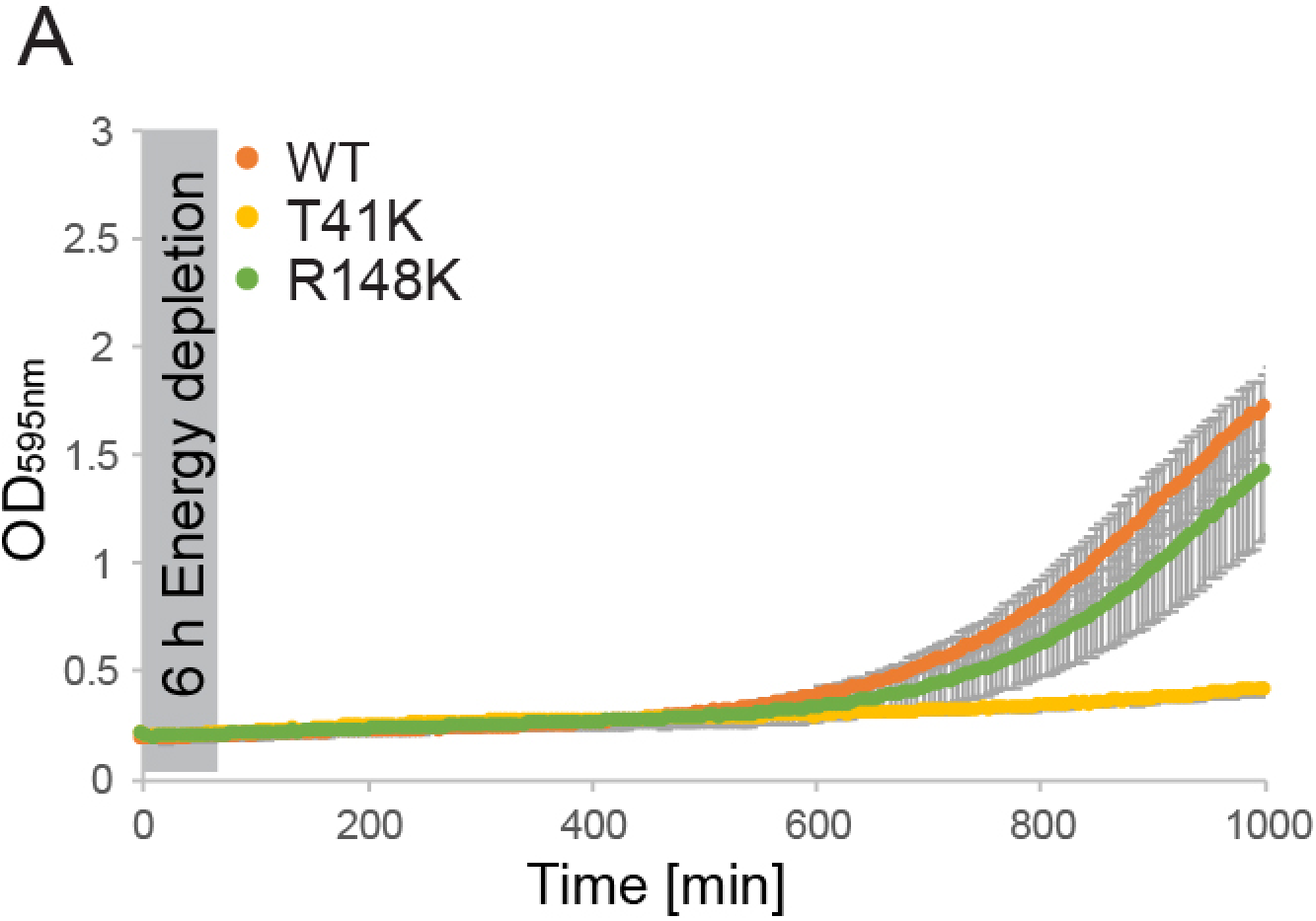
Filament formation is essential for recovery from energy depletion. (A) Growth curves of WT and mutant (T41K, R148K) strains diluted in SC medium to OD 0.05 after 6 hours of energy depletion. Growth was determined by measuring the optical density at 595 nm. Plots were generated from triplicates of 3 different biological replicates and the experiment was carried out at least 3 times.

## Acknowledgements

We thank C. Desroche Altamirano, S. Maharana, J. Guillén Boixet, D. Richter, T. M. Franzmann, C. Iserman and A. Esslinger of the MPI-CBG for critical reading of the manuscript. We are grateful to Prof. M. P. Ashe and S. G. Campbell for sending eIF2B mutant strains that helped us identify the mutations studied in this work. We thank S. Kroschwald for providing the control strain used for Figure S1E. We thank the following members of Services and Facilities of the Max Planck Institute of Molecular Cell Biology and Genetics (MPI-CBG) for their support: B. Borgonovo and R. Lemaitre (Protein Expression Purification and Characterization) for help with protein expression and purification; B. Nitzsche (Light Microscopy Facility) for expert technical assistance regarding SIM; I. Nuesslein, J. Jarrells and C. Eugster Oegema (Cell Technologies Group) for acquisition of FACS data; M. Stoeter (Technology Development Studio) for help performing the *in vitro* reconstitution screen.

We gratefully acknowledge funding from the Max-Planck Society for all authors; additional funding came from the following sources: EN and SA were supported by a grant of the German Research Foundation (DFG, AL 1061/5–1); GM was supported by a DIGS-BB doctoral fellowship; TMF had support from MaxSynBio consortium; and SA had additional funding from Human Frontiers Program grant RGP0034/2017 and Volkswagen “Life?” initiative.

## Author contributions

EN, Conception and design, Acquisition of data, Analysis and interpretation of data, Drafting or revising the article; GM, Conception and design, Interpretation of data; DR, WL, AB, Acquisition of data, Analysis and interpretation of data; TMF, Conception and design, Analysis and interpretation of data; GP and SA, Conception and design, Analysis and interpretation of data, Drafting or revising the article.

---

| Name/Modification | Genotype | Source/Reference |
| --- | --- | --- |
| W303 ADE+ | *MATalpha can1-100 his3-11,15 leu2-3,112 trp1-1 ura3-1* | Gift, Lindquist lab |
|  | **Alternative name** |  |
| W303 ADE+ GCD1::mCherry-SpHIS5 | 3663 | This study |
| W303 ADE+ GCD2::mCherry-SpHIS5 | 3665 | This study |
| W303 ADE+ GCN3::mCherry-SpHIS5 | 3671 | Petrovska et al. |
| W303 ADE+ GCD6::mCherry-SpHIS5 | 3667 | This study |
| W303 ADE+ GCD7::mCherry-SpHIS5 | 3669 | This study |
| W303 ADE+ SUI2::mCherry-SpHIS5 | 4324 | This study |
| W303 ADE+ GCD1::sfGFP(V206R)- KanMX | 3661 | This study |
| W303 ADE+ GCD2::sfGFP(V206R)- KanMX | 3135 | This study |
| W303 ADE+ GCN3::sfGFP(V206R)- KanMX | 3141 | Petrovska et al. |
| W303 ADE+ GCD6::sfGFP(V206R)- KanMX | 3137 | This study |
| W303 ADE+ GCD7::sfGFP(V206R)- KanMX | 3138 | This study |
| W303 ADE+ SUI2::sfGFP(V206R)-KanMX | 3136 | This study |
| W303 ADE+ GCN3::HA-SpHIS5 | 4057 | This study |
| W303 ADE+ *gcn3-Δ* | 3682 | This study |
| W303 ADE+ *gcn3-Δ Gcd1::sfGFP(V206R)-SpHIS5* | 3682 | This study |
| W303 ADE+ *gcn2-Δ* | 3680 | This study |
| W303 ADE+ *gcn2-Δ Gcn3::sfGFP(V206R)-SpHIS5* | 4127 | This study |
| W303 ADE+ *hsp104-Δ* | 3401 | Gift, Lindquist lab |
| W303 ADE+ *hsp104-Δ Gcn3::sfGFP(V206R)-SpHIS5* | 4325 | This study |
| W303 ADE+ GCN3(T41K) | 4288 | This study |
| W303 ADE+ GCN3(R148K) | 4291 | This study |
| W303 ADE+ *trp::GPD-ypHluorin2 GCN3::sfGFP(V206R)-SpHIS5* | 3923 | Munder et al. |
| W303 ADE+ *trp::GPD-ypHluorin2 GCN3(T41K)::sfGFP(V206R)-SpHIS5* | 4275 | This study |
| W303 ADE+ *trp::GPD-ypHluorin2 GCN3(R148K)::sfGFP(V206R)-SpHIS5* | 4279 | This study |
| W303 ADE+ GCN3(T41K)::sfGFP(V206R)-SpHIS5 | 4269 | This study |
| W303 ADE+ GCN3(R148K)::sfGFP(V206R)-SpHIS5 | 4270 | This study |
| BY4741 [RNQ+], pAG415GPD-Rnq1LC-mCherry | 9, O-1354 | Kroschwald et al. |

List of plasmids

List of plasmids
| **Protein expression** | **Alternative name** |
| --- | --- |
| pBIG-GCN3-GCD7-GCD2 | 3031 |
| pBIG-HIS6-GCD6- GCD1 | 3032 |
| **C-terminal tagging** |  |
| pKT128-sfGFP(V206K) | L-264 |
| pKT127-sfGFP(V206K) | L-263 |
| pKT128-mCherry | L-295 |
| pYM45 | L-353 |
| **Integration of pHluorin2** |  |
| pAG304GPD-pHluorin2 | O-3192 |
| **Gene deletion** |  |
| pUG6 | L-178 |
| pUG27 | L-179 |

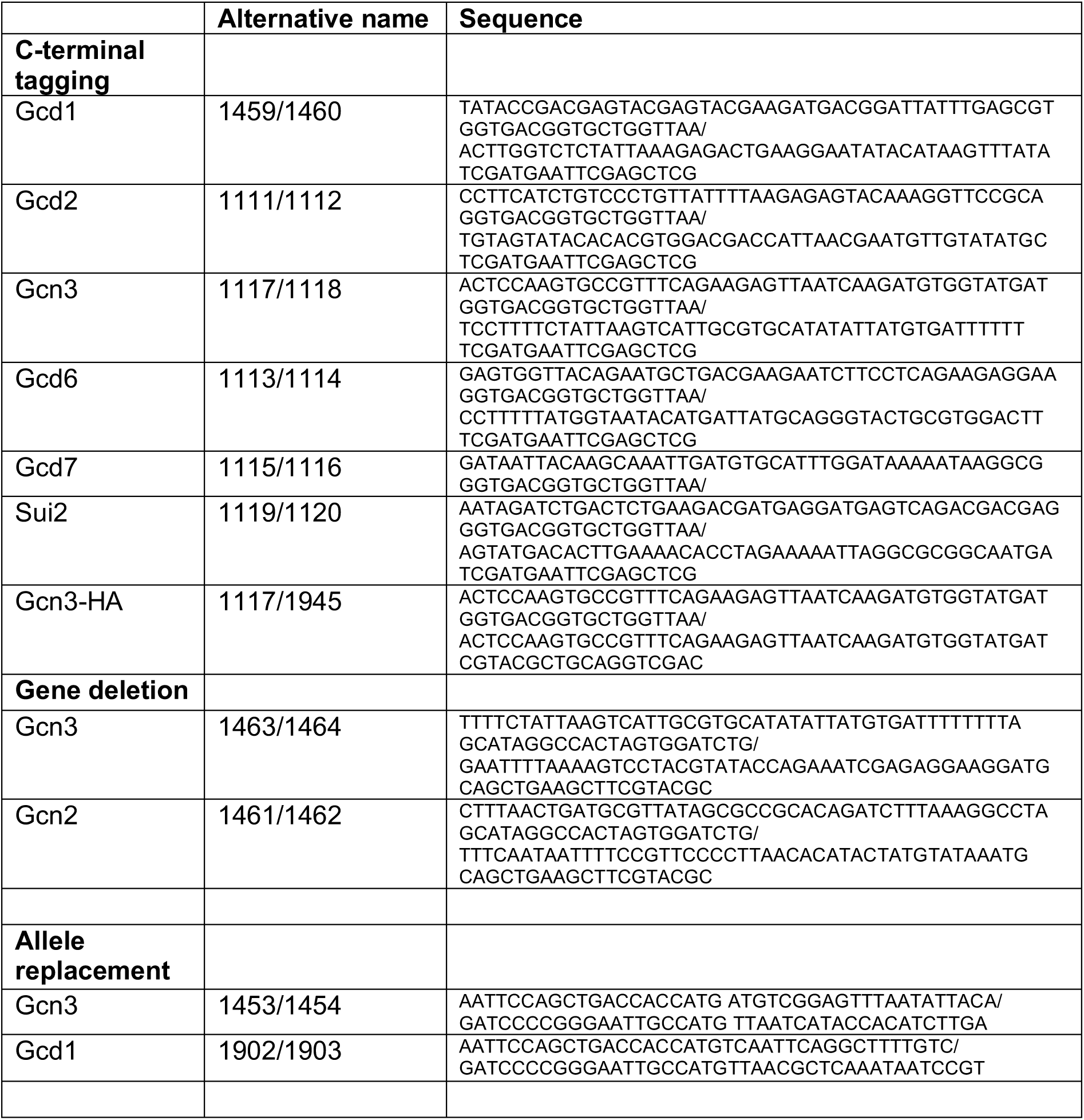
List of primers

